# BetaScan2: Standardized statistics to detect balancing selection utilizing substitution data

**DOI:** 10.1101/497255

**Authors:** Katherine M. Siewert, Benjamin F. Voight

## Abstract

The recently reported statistic *β* detects balanced haplotypes without the need for genomic data from an outgroup species. Here we present an extension to the method that incorporates between-species substitution data into the *β* statistic framework. We show that this approach outperforms existing summary statistics in simulations. We also present the variance of *β* with and without substitution data, allowing calculation of a standardized score. Besides providing a measure of significance, this enables a proper comparison of *β* values across varying underlying parameters, a feature lacking from some related methods.

---

If multiple alleles at a locus are advantageous, then balancing selection can maintain this diversity for long periods of time (Charlesworth, 2006). Statistics designed to detect balanced loci search for the resulting deviations in the patterns of genetic variation (Tajima, 1989; Hudson *et al*., 1987).

The previously developed method *β* identifies a specific signature of long-term balancing selection: the build-up of polymorphisms at near-identical frequencies to one another (Siewert and Voight, 2017). *β* is the difference between two estimators of the mutation rate 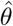, one sensitive to the effects of balancing selection 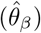 and one that is not (Watterson’s estimator 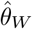). There are two versions of *β*: *β*^(1)^ incorporates outgroup sequence data when available, while *β*^(1)^* only requires a folded site frequency spectrum.

We report two improvements to *β*. The first is a new estimator based on the number of fixed differences with an outgroup species (i.e., substitutions), 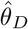. Incorporating this estimator into our *β* framework increases power, owing to the expected reduction of substitutions under balancing selection (Charlesworth, 2006; Hudson *et al*., 1987). Second, we derive the variance of our statistics, enabling standardization of *β*.

We first derive a closed-form solution for the expected number of substitutions under the neutral model **(Supplementary Material)**. This expression leads to an estimator of the mutation rate 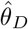 based upon the number of substitutions *S*:

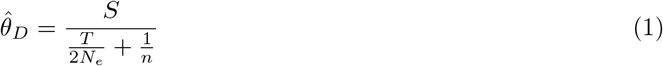

Here, *T* is the divergence time between the two species, *N_e_* is the effective population size, and *n* is the number of surveyed chromosomes. To incorporate information from substitutions, we replace 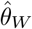 from the original unfolded statistic with 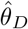 to define *β*^(2)^:

**Figure 1:**
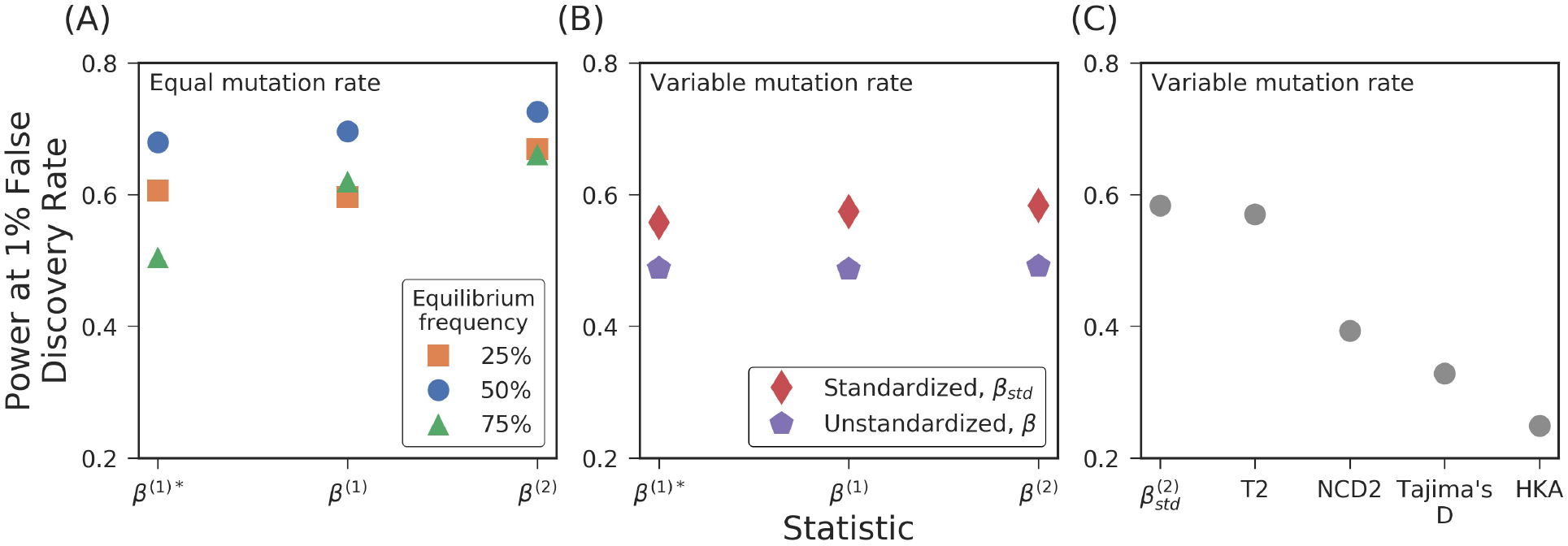
Power of *β* statistics at (A) different equilibrium frequencies and (B) with mutation rate variation, where one half of neutral and balanced simulation replicates had a mutation rate of 2.5 × 10^−8^ (our default rate), and the remaining half had a rate of 1.25 × 10^−8^. (C) Power of *β*^(2)^ compared to other methods for detecting balancing selection. An equilibrium frequency of 50% was used for (B) and (C). The values of each statistic were compared between simulations containing only neutral variants (True Negatives) or with a balanced variant (True Positives).

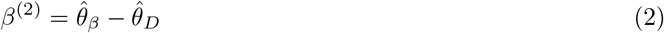

Under long-term balancing selection, variants nearby the selected site are maintained at similar allele frequencies rather than fixing in the population, resulting in an increased estimate of *θ_β_* and reduced estimate of *θ_D_*. Therefore, *β* is expected to score substantially above zero in the presence of long-term balancing selection.

Intuitively, the variances of the *β* statistics depend on the population size, survey sample size, equilibrium frequencies, and mutation rate. We derive the variances of *β*^(1)^* and *β*^(2)^ **(Supplementary Material)**, allowing for comparison of scores across samples and genomic loci where these factors may differ. Our expressions for variance match simulation results **(Supplementary Fig. 1 and 2).** To obtain the variance of *β*^(1)^ we used the framework reported in Achaz (2009) **(Supplementary Material)**. The expected value of all *β* statistics is zero, leading to:

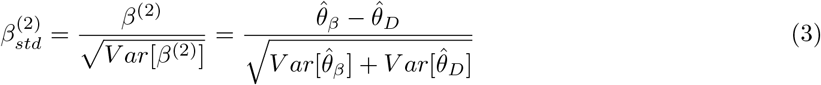

Full derivations for 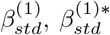, and 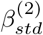 can be found in the **Supplementary Material**. Calculating the variance of all three *β* statistics requires an estimate of 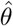, the underlying mutation rate. The variance of *β*^(2)^ also requires an estimate of the speciation time. We discuss techniques for estimation of these parameters in the **Supplementary Material**. However, the *β* statistics are robust to some specification error **(Supplementary Fig. 5 and 6)**. We recommend 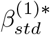 in cases without outgroup data to polarize ancestral states, 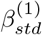 when ancestral states are known but no informative outgroup is available to call substitutions, and 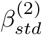 when substitutions are available.

We measure the power of our statistics using simulations **(Supplementary Material)**. We find that *β*^(2)^ has higher power than either *β*^(1)^ statistic, demonstrating that substitution counts provide additional signal over polymorphism data **(Fig. 1a, Supplementary Fig. 3 and 4)**. When there is mutation rate variation across simulations, we find that standardization improves power (**Fig. 1b**).

Next, we compare power to alternative methods: *NCD2* (Bitarello *et al*., 2018), *NCD_mid_* (Cheng and DeGiorgio, 2018), *T*1 and *T*2 (DeGiorgio *et al*., 2014), Tajima’s D (Tajima, 1989) and the HKA test (Hudson *et al*., 1987). When there is mutation rate variation, we find that 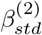 performs the strongest **(Fig. 1c)**. However, *T*2, a method that relies on grids of simulations to generate composite likelihoods, performs best when the mutation rate is stable and the parameters underlying the simulations for *T*2 match the selection scenario. When there is a mismatch between these, the power of *T*2 is reduced, and *β* starts to outperform **(Supplementary Fig. 3 and 4)**. Our results were not biased by window size **(Supplementary Fig. 7)** or power comparison method **(Supplementary Material)**. These analyses show that the *β* framework is a flexible and effective method to detect balancing selection.

Software to calculate all versions of *β* is available at https://github.com/ksiewert/BetaScan.

